# Denoising of Aligned Genomic Data

**DOI:** 10.1101/590372

**Authors:** Irena Fischer-Hwang, Idoia Ochoa, Tsachy Weissman, Mikel Hernaez

## Abstract

Noise in genomic sequencing data is known to have effects on various stages of genomic data analysis pipelines. Variant identification is an important step of many of these pipelines, and is increasingly being used in clinical settings to aid medical practices. We propose a denoising method, dubbed SAMDUDE, which operates on aligned genomic data in order to improve variant calling performance. Denoising human data with SAMDUDE resulted in improved variant identification in both individual chromosome as well as whole genome sequencing (WGS) data sets. In the WGS data set, denoising led to identification of almost 2,000 additional true variants, and elimination of over 1,500 erroneously identified variants. In contrast, we found that denoising with other state-of-the-art denoisers significantly worsens variant calling performance. SAMDUDE is written in Python and is freely available at https://github.com/ihwang/SAMDUDE.

## Introduction

Sequencing data have expanded our understanding of genetic material and their role in biological processes, opened up new areas of biological inquiry, and are guiding the trajectory of modern biomedical research^1^. Raw sequencing data are stored in the FASTQ file format and converted to the SAM file format following alignment to a reference genome. Both file types comprise sequences of nucleotide bases called “reads”, which are accompanied by sequences of quality scores that indicate the sequencing machine’s confidence in the base calls making up the reads. However, the genomic sequencing process is imperfect and can result in reads containing various types of noise including base substitutions, insertions, and deletions (INDELs).

Although noise characteristics vary across sequencing technologies, they are well characterized for some sequencing platforms. For example, Illumina sequencing technology produces “short” reads on the order of hundreds of bases, with an average substitution error rate of less than 1%, and insertion and deletion rates orders of magnitude lower^2^. Furthermore, these errors were found to be correlated with position within the read, resulting in position-dependent noise characteristics. These errors can affect downstream applications, with an important application being variant calling, or the identification of genetic polymorphisms unique to individuals. Variant identification from WGS is increasingly being used to diagnose, gain biological insight to, and design treatments in the clinical setting, especially in the field of rare genetic disease research^3^. Thus, accuracy of variant identification is paramount.

Algorithms for removing noise, or denoisers, have been proposed for not only genomic sequencing data^4^, but also for other biological methods relying on genomic sequencing, like ChiP-seq^5^. Denoisers for genomic sequencing data change individual bases in reads while retaining the original quality scores. They are typically tested on simulated and real data sets in FASTQ format, and have been shown to perform well on some of the early stages of genomic sequencing pipelines, such as correcting base calling errors in the simulated data sets, increasing both breadth and depth of reads coverage during alignment^6^, or improving de novo assembly of real data sets^7^. However, these analyses often do not extend to later steps in genomic sequence analysis pipelines, and those that do focus on non-human data sets^8^. To our knowledge, none of these works examines the effect that denoising might have on variant calling. The variant calling procedure is complex and relies on alignment information, including quality scores which are a direct function of the analog signals used to determine the called base. In fact, a survey of lossy quality score compressors has already shown that changing quality scores alone can sometimes have a beneficial effect on variant calling^9^. This result shows that in a sense, lossy quality score compression denoises the genomic data, resulting in more accurate variant calling. Taken together, the existing body of work on denoising and variant calling suggests that read denoising procedures that leave quality scores unchanged may result in unexpected variant calls, and that effective read denoising must be accompanied by quality score updates.

In this work, we propose a novel denoising method, SAMDUDE, which takes advantage of alignment information contained in the SAM file in order to both denoise reads and update quality scores. We evaluate the effect of denoising on variant calling by comparing variants identified in files before and after denoising by SAMDUDE. We also evaluate files that have been denoised using other state-of-the-art denoisers that operate solely on reads in FASTQ files. This variant calling comparison methodology has already been used to analyze the effect of lossy compression on quality scores beyond the early steps in a genomic sequencing pipeline^9^. To our knowledge, this is the first application of such a comparison methodology on denoised genomic sequencing data, and provides a unique framework for directly evaluating the effect of denoising on sequencing data. To highlight the potential utility of simultaneous base denoising and quality score updating in a clinical setting, we perform denoising and variant calling comparisons on human data sets. We show that the simultaneous reads denoising and quality score updating procedure either maintains or improves variant calling with respect to the original SAM file, while denoising schemes that change only the reads result in degraded variant calling performance.

### Survey of denoisers for genomic data

Current state-of-the-art denoisers perform denoising based on a variety of techniques including *k*-mer counting and statistical error models, and target either substitution errors, insertion and deletion errors, or a combination of both^4^. We chose Musket^10^ and RACER^11^ to serve as benchmarks for SAMDUDE denoising performance, since both were touted for their ability to handle human WGS sequencing data sets^6^. In addition to Musket and RACER, BFCounter^12^ and Lighter^13^ are also preferred in the field for their memory efficiency and speed, respectively, and are thus the most likely to be used in practice. We omitted comparisons with BLESS 2^14^, another denoising tool especially popular for its speed, due to installation difficulties. Here, we briefly describe the techniques underlying these four denoisers.

Musket^10^ uses a *k*-mer spectrum approach in which reads that are suspected to be erroneous are changed until their *k*-mers appear frequently in the entire data set. The *k*-mer spectrum is constructed using a parallelized master-slave model, resulting in Musket’s highly competitive execution time and excellent parallel scalability. Denoising is performed using a multistage workflow which begins with multiple iterations of two-sided conservative base correction. Two-sided conservative base correction is followed by multiple iterations of one-sided aggressive correction and voting-based refinement. Musket is able to denoise paired-end reads data sets simultaneously.

RACER^11^ uses a *k*-mer counting approach to denoise FASTA and FASTQ data. Its *k*-mer counting method retains *k*-mers with counts above a given threshold, while correcting all other ones. RACER utilizes a unique and efficient hash table-based data structure which makes it extremely space efficient. While RACER does not denoise both files in a paired-end reads data set at the same time, each of the FASTQ files can be denoised independently and recombined in subsequent analysis steps. RACER requires approximate genome size as a parameter.

BFCounter^12^ also uses a *k*-mer counting approach coupled with a Bloom filter in a two-pass denoising process. The use of a Bloom filter results in reduced memory requirements of nearly 50% memory savings, as compared to popular *k*-mer counting software. However, the two-pass implementation requires a significant amount of time for completing denoising, especially on human WGS data. Like RACER, BFCounter requires approximate genome size as a parameter.

Unlike the previously mentioned denoisers, Lighter^13^ avoids *k*-mer counting and instead relies entirely on Bloom filters to perform denoising. Compared with most denoising methods, Lighter is extremely fast and memory-efficient, but like RACER and BFCounter it also requires an estimate of genome size as a parameter.

## Results

To formulate the proposed denoising method, we assume a setting in which a genetic sample undergoes high-throughput shotgun sequencing, producing a large number of short, overlapping reads of length on the order of hundreds of base pairs. The errors introduced during the sequencing process are assumed to be primarily substitution errors, while INDELs are assumed to be negligible. We also assume that a reference genome is available, and that the reads can be aligned to the reference.

Our proposed denoising method, SAMDUDE, is based on the Discrete Universal Denoiser (DUDE) algorithm proposed in^15^. DUDE is a sliding-window discrete denoising scheme which is universally optimal in the limit of input sequence length when applied to an unknown source with finite alphabet size corrupted by a known discrete memoryless channel. The universal optimality of the DUDE guarantees that in the asymptotic limit of input sequence length it does as well as the best scheme of its type, regardless of the characteristics of the underlying noise-free sequence. In brief: DUDE uses a two-pass procedure to first infer statistics of the source sequence based on the noisy sequence, and to then denoise the noisy sequence using the inferred statistics and the noise channel characteristics. The DUDE setting and algorithm is illsutrated in Figure 1.

**Figure 1.**
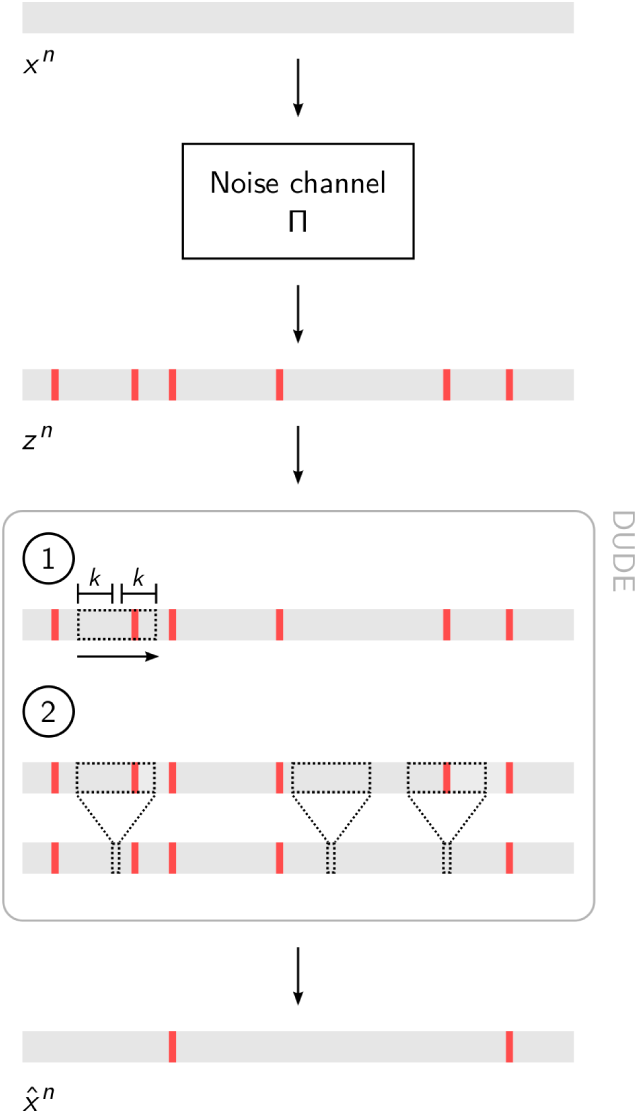
Schematic of the DUDE setting and algorithm. A noise-free sequence, *x*^*n*^ (gray bar), is corrupted by a noise channel **Π**, producing noisy sequence *z*^*n*^ with errors (red lines). In the first pass of DUDE, a sliding window (dashed rectangle) obtains vectors of counts for contexts of length 2*k*. In the second pass, the counts vectors of each context, together with the channel statistics, are used to denoise the central symbol of the context (small dashed rectangles).

In order to apply the DUDE-like denoising framework to the genomic sequencing setting, we make certain algorithm design choices based on assumptions about the problem setting. While the universal denoising setting assumes a single noise-free input sequence and a single noise-corrupted output sequence of equal length, in the high-throughput sequencing setting the channel input is a single, noise-free sequence and the output are numerous overlapping, short, noisy sequences. The reads may not necessarily all be of the same length, but are all assumed to be much shorter than the noise-free sequence length. Despite the difference between these settings, the sequencing reads can be thought of as samples from a single noisy sequence that can be inferred from the reads using alignment information. Under this assumption, we aggregate information from each read into statistics about the inferred noisy sequence. The universal denoising setting also assumes that the noise channel is memoryless and known, and corrupts sequences only with substitution errors. The error characteristics of sequencing technologies are generally known, but in order to account for individual variations in performance from machine to machine and to avoid the potentially confounding effects of difficult-to-sequence regions in the genome, we use alignment information in the SAM file to generate our own estimate of the channel’s noise-injecting characteristics. We consider only the main mapping of the aligned reads, disregarding any other mappings. Finally, while paired-end reads are acceptable inputs to the denoiser, the pairing information is not used in the denoising process. The SAMDUDE denoising scheme is described in detail in the Methods section.

While all sequencing technologies inject all three types of errors, the noise channel model used in SAMDUDE is a particularly accurate reflection of Illumina sequencing technologies. Furthermore, due to the importance of variant identification in the clinical setting, we evaluated the effect of denoising on variant calling in human data sets. We tested SAMDUDE on three different paired-end WGS data sets of the *H. Sapiens* individual NA12878. The data sets are: ERR262997 corresponding to 30× -coverage, CEUTrio.HiSeq.WGS corresponding to 100× -coverage, and NA12878_V2.5_Robot_2 corresponding to 40× -coverage. For convenience, we refer to these data sets as 1, 2 and 3, respectively. Variant calls were compared against the version 37 gold standard call set for individual NA12878 released by the National Institute of Standards and Technology’s (NIST) Genome in a Bottle consortium (GIAB)^16^. While the GIAB gold standard call set is a fairly conservative estimate of the individual NA12878 true variant call set, it is widely regarded by the field as the standard benchmark for evaluating sequence analysis algorithms. Furthermore, there exist “gold standard” (consensus of polymorphisms) variant call sets for certain human individuals, which can be used for an intuitive and direct method of assessing variant calling performance.

Denoising performance was evaluated with respect to variant calling of single nucleotide polymorphisms (SNPs). We first analyzed the effect of denoising on variant calling performance for individual human chromosomes using SAMDUDE and other state-of-the-art denoisers. We also compared the effect of denoising reads to the effect of lossy quality score compression. Lossy quality score compressors were developed to decrease the size of SAM files while still maintaining variant calling performance. Previous work showed that while quality score compressors effectively reduced SAM file size, they also sometimes had the unintended effect of improving variant calling performance^9^. For this reason, the lossy quality score compressors serve as a counterpoint to the SAMDUDE algorithm’s procedure of changing both reads and quality scores in tandem with the explicit goal of improving variant calling performance. Finally, we analyzed the effect of SAMDUDE denoising on human WGS data.

### Human chromosome denoising with SAMDUDE

For individual chromosome denoising experiments, we chose to use chromosomes 11 and 20. Chromosome 11 was chosen as representative of the median chromosome length in the human genome, and chromosome 20 was chosen since it is frequently used in genomic data tool assessment as representative of a small human chromosome^9^. The leftmost column of Figure 2 shows that SAMDUDE can have varied effects across different data sets. For data set 1, denoising with SAMDUDE resulted in an increase in T.P. variants called concomitant with a decrease in F.P. variants, resulting in a modest gain in F-score. Although denoising resulted in a slight decrease of T.P. variants called in data set 2 relative to the original file, it also resulted in a very large decrease in number of F.P. variants called, resulting in a large gain in F-score. In contrast, a handful of additional T.P. and F.P. variants were called in data set 3, resulting in no change in F-score. Despite variation in results across data sets, the effect of SAMDUDE denoising is consistent across chromosomes within a given data set.

**Figure 2.**
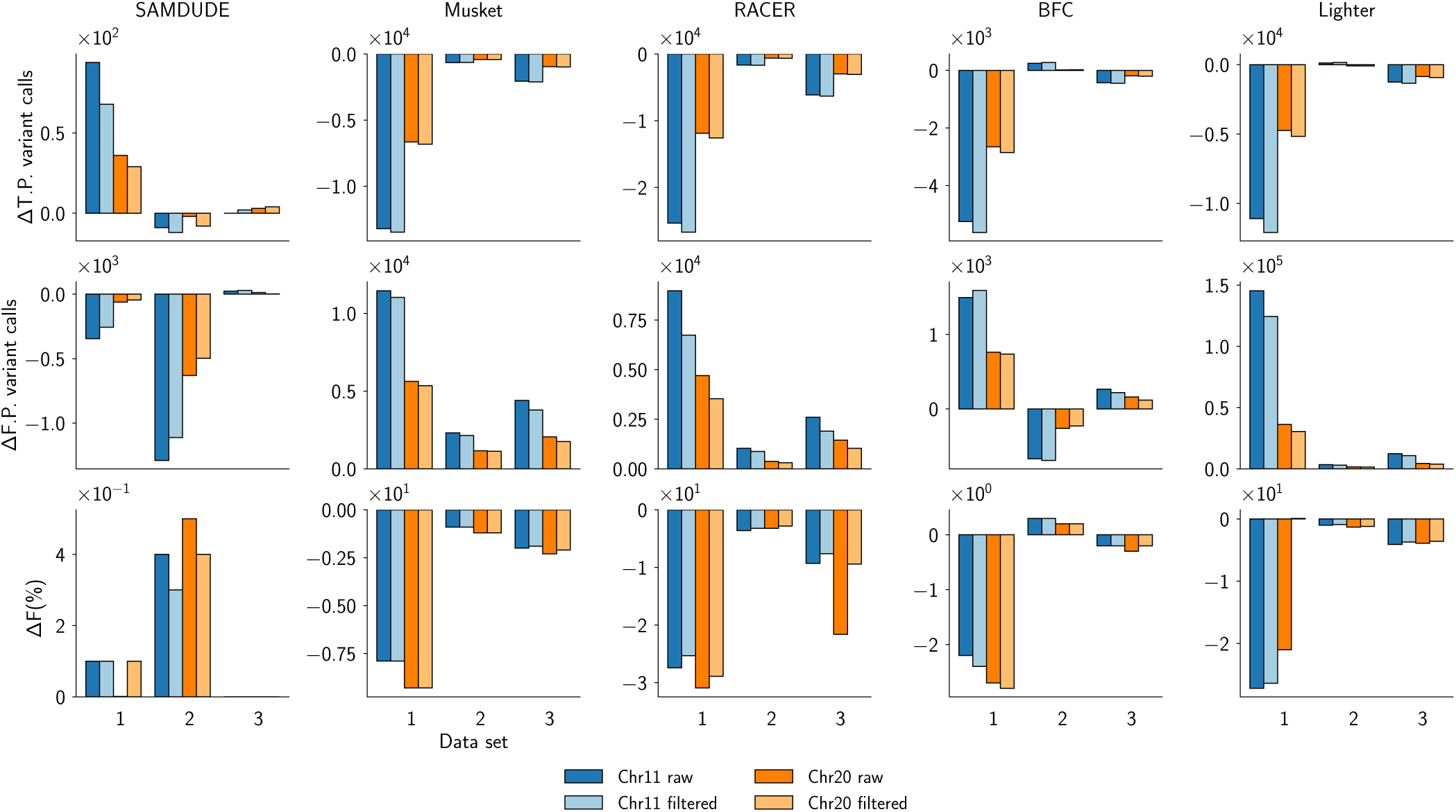
Denoising results for SAMDUDE (left column), Musket (second from left), RACER (center), BFCounter (second from right), and Lighter (right) for chromosomes 11 (blue) and 20 (orange). Positive changes in T.P. and F indicate increases in T.P. variants called and improvement of F-score, respectively. A negative change in F.P. indicates that fewer variants were erroneously called. Raw (dark colors) and filtered (light colors) variant call values are shown.

While differences in performance across data sets seem to indicate inconsistency in the SAMDUDE denoising algorithm, these results make sense in the context of coverage and initial data quality, which are summarized in Figure 3. The original performance metrics for data set 3 are by far the highest. In contrast, while data set 2 also has a high initial sensitivity, its relatively low precision leaves room for improvement in F-score. Data set 1 has the most room for improvement, with low initial sensitivity, precision and F-score.

**Figure 3.**
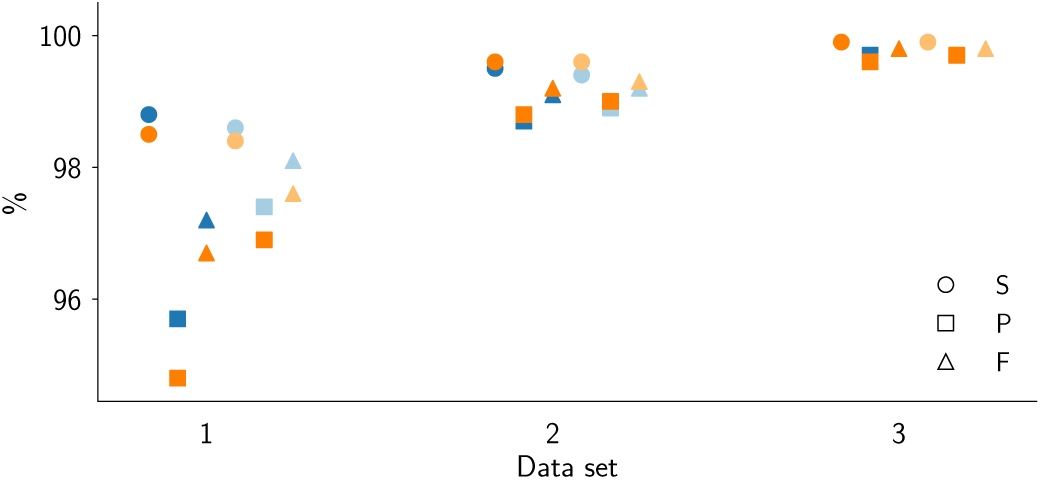
Statistics for variant call sets generated from the original data sets (blue, chromosome 11 and orange, chromosome 20). The statistics are: sensitivity (circles), precision (squares) and F-score (triangles). Statistics are grouped by variant call type: raw (dark colors) and filtered (light colors).

Because of SAMDUDE’s conservative behavior, it might be suspected that the gains from updating quality scores are small compared to the effect of denoising bases in the reads. To test this hypothesis, we created partially-denoised SAM files where the reads were those obtained from SAMDUDE denoising, but were paired with the original quality score strings. These partially-denoised files were then analyzed using the same variant calling pipeline, and the results of this experiment are summarized in Supplementary Table 1 (labeled Partial Denoising). For all data sets, partial denoising resulted in either no change or an increase in F-score, even after GATK filtering. However, for data set 1 and 2 the gains were not as large as those attained using SAMDUDE after GATK filtering of variant calls.

**Table 1.** Results for raw and filtered variant calling on denoised data set 1 using SAMDUDE and Musket.

|  | T.P. raw | T.P. filtered | F.P. raw | F.P. filtered |
| --- | --- | --- | --- | --- |
| SAMDUDE | 1,954 | 1642 | -1,509 | -1,923 |
| Musket | -208,144 | -215,524 | 324,957 | 305,362 |

It might also be suspected that the same amount of random base changes would also result in improvements in variant calling. To check this hypothesis, we ran the variant calling pipeline on the SAM files for chromosome 20 with bases changed at random based on a uniform distribution over the possible nucleotide bases (see Supplementary Table 2). The results of this experiment are summarized in Supplementary Table 1 (labeled Random noise). For all data sets, addition of random noise resulted in either no change or a decrease in sensitivity, concurrent with a uniform increase in precision. Overall, the effect of random noise is an increase in F-score for all raw variant calls, but a decrease in F-score for GATK-filtered variant calls in data sets 1 and 2.

Together, these results support our claim that SAMDUDE is a conservative denoising method which will not adversely affect data sets that do not need denoising, while improving those that can benefit from denoising. Furthermore, SAMDUDE’s performance is robust and consistent for both raw and filtered variant calls, unlike that of partial denoising and random noise addition.

### Comparison of SAMDUDE denoising with state-of-the-art genomic denoisers and quality score compressors

The effects of denoising by the state-of-the-art genomic denoisers Musket, RACER, BFCounter and Lighter are shown in the center and right columns of Figure 2. In most cases, the denoisers resulted in significant decreases in the number of T.P. variants called, with significant increases in the number of F.P. variants called, leading overall to significant decreases in F-score for both the raw and GATK-filtered variants. The only exception is the denoising of data set 2 using BFCounter, which resulted in a slight increase in T.P. variants called, slight decrease in F.P. variant called, and a corresponding slight increase in F-score. However, this improvement in variant calling is not consistent, as denoising data sets 1 and 3 using BFCounter resulted in worse F-scores. These changes show that although current state-of-the-art denoisers have been shown to improve early steps of the genome analysis pipeline, their denoising choices tend to have adverse effects on variant calling.

The trends are consistent even when we consider different variant call filtering levels. Supplementary Figures 1-6 show the variant calling precision as a function of sensitivity for different filtering criteria. We focus on the variant call filtering results for data set 2 (Supplementary Figures 3 and 4) since for this data set SAMDUDE denoising resulted in the largest number of total changes to the variant call set, and also in the largest change in F-score. We also focus on comparing sensitivity filtering for the results of Musket and RACER due to their consistent denoising trends. To construct these curves, we set percentile thresholds starting at the 10^th^ and ending at the 90^th^ percentile at increments of 10% for quality of depth (QD) of the raw variant calls. All variants with QD below the threshold were filtered out, and the remaining variant calls were evaluated against the gold standard call set. The top row of Supplementary Figure 4 shows that even under these filtering criteria, the curves corresponding to the variant call sets of Musket- and RACER-denoised SAM files lie far below the others. In other words, for a given sensitivity level, the variants called under Musket- and RACER-denoised SAM files have significantly worse precision than SAMDUDE.

**Figure 4.**
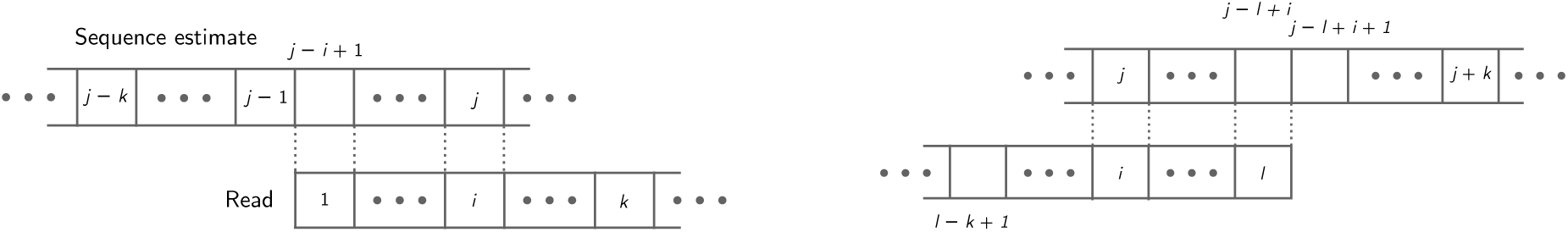
Reads are padded with bases taken from the sequence estimate during the context acquisition and denoising processes. **(a)** Position *i* in the read corresponds to position *j* in the sequence estimate, and 1 ≤ *i* ≤ *k*. The left half of the context for that base is padded with sequence estimate bases from positions *j* - *k* up to *j* - *i* + 1. **(b)** Here, *i* ≥ *l* - *k* + 1, where *l* is the length of the processed read. The base at position *i* can now use positions *j* - *l* + *i* + 1 up to *j* + *k* of the sequence estimate as a right-hand context.

In addition to variant call filtering results for call sets from SAMDUDE, Musket-, and RACER-denoised SAM files, the bottom rows of Supplementary Figures 1-6 also show the filtered variant call set curves for the original call set, the call set resulting from the addition of random noise to the reads, and two call sets resulting from lossy compression of the original file using P-Block and R-Block with compression parameters 3 and 40, respectively. P-Block and R-Block are two state-of-the-art lossy quality score compressors^17^ which have also been shown to improve variant calling performance. Another state-of-the-art lossy compressor, QVZ^18^, has been omitted from the analysis since it is unable to compress SAM files with variable length reads. Again we focus our discussion on data set 2 and observe that in Supplementary Figure 4 shows that the effect of lossy compression on the variant call set is almost indistinguishable from that of SAMDUDE. However, the rightmost points in each of the filtered variant call set curves show that SAMDUDE outperforms all other schemes when sensitivity and precision are both high. SAMDUDE’s dominance, especially at high sensitivity levels for chromosome 11, makes sense since SAMDUDE’s denoising performance improves as read coverage increases.

### Human whole-genome denoising with SAMDUDE

Next, we evaluated the effect of denoising on variant calling for an entire WGS human data set. For this experiment, we used all reads of data set 1, and the results are summarized in Table 1. After SAMDUDE denoising the raw variant call set included in total 1, 954 additional true positive variants. Furthermore, 1, 509 false positive variants were eliminated relative to the original set of variant calls. After the GATK pipeline, 1, 642 of the additional true positive variants were validated, and the number of additional false positive variant calls eliminated increased to 1, 923. In contrast, raw variant calls based on the Musket-denoised file resulted in 208, 144 fewer true positive calls and 324, 957 additional false positive calls relative to the original set of variant calls. After the GATK pipeline, the number of true positive variant calls missed increased to 215, 524 while the number of false positive variant calls increased to 305, 362. Variant calls from the RACER-denoised data set were unable to be validated against the gold standard call set due to pipeline errors.

## Discussion

We have presented SAMDUDE, a denoising method that uses alignment information in SAM files and a statistical model of the genomic data in order to improve variant calling. Because of the assumptions used by the denoising model, SAMDUDE’s intended use is for improving the quality of short-reads sequencing reads obtained from healthy tissue.

Taken together with the initial data quality statistics in Figure 3, the range of improvements observed in Figure 2 imply that SAMDUDE is a “conservative” denoising algorithm that makes few or no changes to the reads and quality scores when the original data set is already of very high quality, but makes sound denoising choices resulting in variant calling improvements when the original data set is of lesser quality. The quality of denoising performance also correlates with data coverage: data set 2 has the highest coverage and SAMDUDE denoising performance is the best on this set. Since SAMDUDE relies on empirical estimates of *k* - mer distributions and the noise channel, the higher the coverage and the more accurate the empirical estimates, resulting in better denoising performance. It is also notable that the trends in performance metric changes hold for both raw variant and filtered variant calls, adding to our confidence in SAMDUDE as a conservative denoiser that integrates well with existing recommended genomic data analysis pipelines.

The results of denoising aligned whole genome data are consistent with those observed for individual chromosomes. While the relative number of true positive variants identified might seem relatively small, the extra information provided by each extra variant could be invaluable. Single point mutations are responsible for numerous human diseases, and other diseases once assumed to be caused by a single variant with large effect are now being understood to be the result of multiple monogenic mutations, or of collections of rare variants in previously identified genes^19–21^. Perhaps more importantly, the elimination of false positive variants is crucial to accurate diagnosis and appropriate treatment design^22–24^. Thus, in the clinical context the implications of every additional true positive variant identified and each false positive variant eliminated are far larger than the objective tally.

Our results emphasize that the quality score updating step of SAMDUDE is crucial to improving variant calling outcome, and that denoising reads alone is insufficient for higher quality of variant calls. SAMDUDE was able to both identify thousands of additional variants and eliminate a similar number of false positive variants from a single human whole-genome data set. In contrast, state-of-the-art denoisers, which were designed to improve earlier steps in the sequencing pipeline and are limited to changing information only in the reads and not quality scores, led to degraded variant calling performance. Our results also highlight the importance of evaluating denoisers on the variant calling step of the genomic sequencing pipeline using real data sets with gold standards.

As a proof-of-concept denoiser, SAMDUDE shows great promise in improving the accuracy of variant calling based on individual sequencing data sets. Furthermore, these encouraging results motivate further experimentation of the parameters and elements of the denoising procedure, including context length *k*, majority and confidence thresholds, quality score updating rule, and a additional refinement of the implementation in order to reduce computational memory and time requirements. We anticipate that the SAMDUDE denoising method will result in an efficient and powerful denoising software that will be a valuable tool for researchers and clinicians alike.

## Methods

### Problem setting

We have the following problem setting: *x*^*n*^ is the true genomic sequence of length *n*, and the sequencing procedure generates a set of *m* noisy reads { *z* _(1)_, *z* _(2)_, …, *z* _(*m*)_} with components taking values in the set of all possible nucleotide bases. The reads are accompanied by a set of quality score strings { *q* _(1)_, *q* _(2)_, …, *q* _(*m*)_} with components taking values in the set of ASCII characters quantifying basecalling quality on a quality score scale. The denoiser has access to the reads and the quality score strings, as well as to the alignment information of each of the reads to the reference sequence. Our goal is to both denoise the bases in the reads and to update the corresponding quality scores in order to improve the accuracy of variant identification, while still preserving polymorphisms that are unique to the individual. Note that in this setting, the true genome sequence *x*^*n*^ is unknown.

In the given problem setting, we assume that for a particular location *i* in the reference genome, the majority of reads covering that position will have base calls that agree. The minority of base calls that do not match with the majority are likely to be errors. Under this assumption, the sequence estimate is obtained by recording the majority base, for some majority threshold, for all reference genome positions covered by the reads. The noise statistics are represented by the noise channel estimate 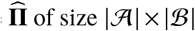. In our setting, the channel input is assumed to be the noise-free sequence taking values in alphabet 𝒜 = {*A, C, G, T*}, while the channel output is a tuple of the called based and the quality score associated with that base. Hence the output alphabet is given by ℬ = 𝒜 × 𝒬, where 𝒬 denotes the alphabet of the quality scores.

The typical size of 𝒬 is 42, which carries a significant computational burden due to the resulting size of 𝒜 × 𝒬. Hence, we adopt the quality score binning method recommended by Illumina for reducing quality score resolution^25^. This method reduces the original alphabet of quality scores from 42 to only 8 bins; hence, 𝒬 ∈ {bin_1_, bin_2_, …, bin_8_}, with bin limits corresponding to those recommended by Illumina (see Supplementary Table 3). We consider this set of output tuples since certain sequencing technologies, like the Illumina technologies, are known to produce reads with position-varying noise characteristics. Typically, different noise characteristics would be characterized by different noise channels. By considering the nucleotide base with its quality score, we can broadly account for various possible position-dependent trends in noise without setting hard boundaries and limiting ourselves to particular assumptions about the noise characteristics. Details on the computation of 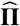 are given in subsection Channel and sequence estimation.

The vector of counts **m** (*l*^*k*^, *r*^*k*^), of size | ℬ|, records the number of times the subsequence *l*^*k*^ *br*^*k*^, comprising left and right contexts *l*^*k*^ and *r*^*k*^, is observed in the collection of reads, with *l*^*k*^, *r*^*k*^ ∈ 𝒜 ^*k*^, and *b* ∈ ℬ. We limit the alphabet of the context to 𝒜 to ensure that each possible context is observed a significant number of times. Details on the computation of the counts vector are given in subsection Counts vector acquisition.

Once the noise channel estimate 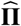 and vectors of counts **m** are acquired, SAMDUDE proceeds as follows. For ease of exposition and with some abuse of notation, we denote an arbitrary read as *z* with *i*^th^ component *z*_*i*_ accompanied by quality score *q*_*i*_. Subsequences of *z* are denoted as 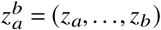.

1. For each base *z*_*i*_ and its associated quality score *q*_*i*_ in read *z*, identify the length 2*k* context string 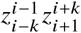 surrounding position *i*, and the bin bin_*i*_ to which *q*_*i*_ belongs.
2. Calculate the estimated probability of observing the left and right contexts 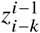 and 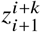 in *x*^*n*^ with different central symbols belonging in 𝒜. The probability is distributed over all symbols in 𝒜 and given by

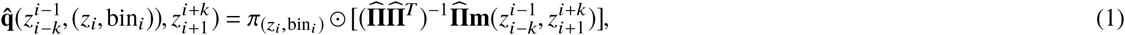

where 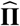 is the channel estimate matrix comprising column vectors {*π*_1_, *π*_2_, …, *π* _|ℬ|_}, and ⊙ represents element-wise multiplication for vectors (see^15^, Eq. (25)).
3. Replace *z*_*i*_ with the base corresponding to the argument of the maximum of the distribution estimate, and update *q*_*i*_ using the maximum of the distribution estimate (see Section Quality score updating for details on the quality score update rule).

We devote the following subsections to detailed descriptions of the components of SAMDUDE: estimates of the channel and true sequence, noise statistics, and vectors of counts for every possible context of size 2*k* observed in the collection of reads. We also describe the quality score updating procedure, specify implementation details and describe the overall evaluation pipeline workflow.

### Channel and sequence estimation

To compute the channel estimate 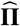, we use alignment information from the SAM file to perform a sequence pileup at every reference genome position by cataloging all reads at that position. At each position we assume that the majority base, for some majority threshold *t*_*m*_ ≥ 0.5, is the true base at that position. That is, the base in the pileup with normalized counts greater than or equal to *t*_*m*_ is declared to be the majority base. If there is no clear majority base, then we do not use information from that position for channel estimation. This rule allows us to use the overwhelming majority of genomic positions in order to estimate the channel characteristics. Additionally, the threshold ensures that information is not used from positions in the genome that display heterozygosity, e.g., due to polyploidy of the organism being sequenced. This, in turn, prevents the conflation of differences in reads overlapping heterozygous positions with noise, and precludes erroneous denoising of reads at those positions. For each base in 𝒜 we record the number of bases in an 4 × 32 conditional counts matrix

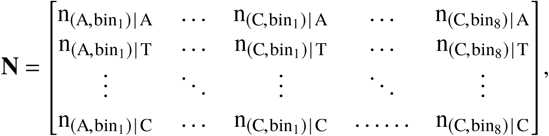

where 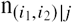 is the number of positions in all reads for which the read contains base *i*_1_ with accompanying quality score in bin *i*_2_ at a position whose majority base is *j*. **N** is row-normalized to obtain 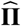

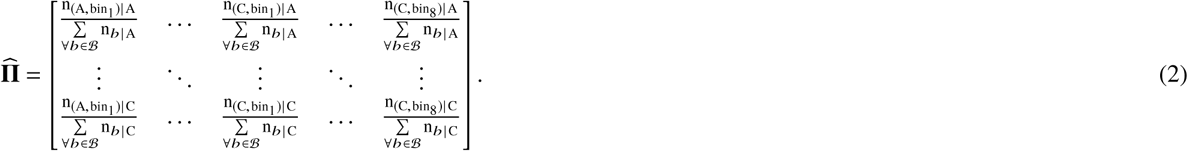

The majority bases are recorded as the sequence estimate.

### Counts vector acquisition

The vectors of counts **m**(*l*^*k*^, *r*^*k*^) record all appearances of context *l*^*k*^*r*^*k*^ surrounding a central symbol *b*, with *l*^*k*^, *r*^*k*^ ∈ 𝒜^*k*^ and *b* ∈ ℬ. For each *b* appearing between the left and right context components *l*^*k*^ and *r*^*k*^, respectively, we record the number of times the sequence *l*^*k*^ *br*^*k*^ appears in the collection of reads as the *b*^th^ component of **m**(*l*^*k*^, *r*^*k*^):

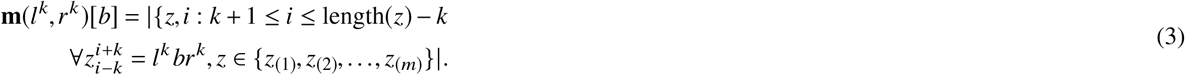

For positions *i* < *k* + 1 or *i* > length(*z*) - *k*, we employ information from the sequence estimate to acquire context counts. This process is detailed in section SAMDUDE implementation details.

### Position-based denoising rules

Denoising is performed at read positions where *z*_*i*_ as well as the surrounding context string contain only bases in 𝒜 = {A, T, G, C}, excluding symbols indicating ambiguity. We use this simple genomic alphabet in order to avoid basing denoising decisions on non-uniquely identifiable context strings. We also set a quality score threshold above which denoising is not attempted. Initial denoising experiments showed that bases changed under SAMDUDE are overwhelmingly those with a low quality score value (see Supplementary Figure 7). Furthermore, we found that bases that have been assigned a high quality score during sequencing are most likely not erroneous and may not benefit from denoising. With these considerations, we established a confidence threshold *t*_*p*_; denoising is not attempted at read positions with quality scores corresponding to a probability above *t*_*p*_.

### Quality score updating

The maximum of the conditional distribution estimate 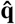 is used to update the quality score accompanying the denoised base. The updating procedure depends on whether the base the denoiser selected matches the original base *z*_*i*_. If the maximum of 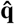 corresponds to the same base as the original one, the quality score is adjusted as follows: convert the original quality score *q*_*i*_ into a confidence probability

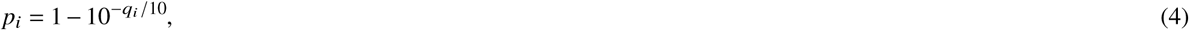

take the arithmetic mean of *p*_*i*_ and the maximum of the estimated conditional probability 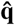, denoted by *p*_max_, and back-convert the averaged probability into the updated quality score. In other words, the updated quality score is

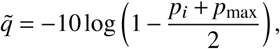

when the denoiser does not recommend a different base. The updated quality score is re-inserted into the quality score string after conversion to an ASCII character as per the sequencing machine’s quality score encoding method. For example, if the sequencing machine encodes on a Phred+33 scale, the quality score string’s *i*^th^ component is replaced with the ASCII character for 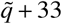. On the other hand, if the denoiser recommends a base change, *q*_*i*_ is simply replaced with 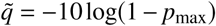. Again, the quality score string’s *i*^th^ component is replaced with the appropriately encoded 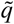.

This procedure was chosen in order to balance the denoiser’s conditional probability estimates with the original quality scores, which reflect the sequencing machine’s confidence in the base calls. Since the original quality score is a function of the original base call, if the denoiser agrees with the basecall, the denoiser’s probability estimate should be combined with the sequencer’s quality score. However, if the denoiser decides on a different base, then the original quality score is unrelated to the denoiser’s chosen base and we can disregard the original quality score in favor of the quality score converted from the denoiser’s probability estimate.

### SAMDUDE implementation details

The reads reported from a sequencing machine cannot always be mapped directly to the reference genome in their entirety, since they may contain bases that are insertions relative to the reference genome, lack bases that correspond to deletions from the reference genome, or contain stretches of bases at the beginning and end of the read that simply do not match the reference genome. These inconsistencies relative to the reference genome are summarized by the sequence aligner in a CIGAR string accompanying the read (https://samtools.github.io/hts-specs/SAMv1.eps). Additionally, large portions of the read may be assigned very low quality scores, indicating entire regions of the read for which the sequencer has low confidence. One strategy for dealing with these inconsistencies is to simply eliminate non-matching or low-quality bases, but this can lead to loss of potentially valuable information. Instead, we retain this information and tailor our use of it to process the reads during channel estimation, counts vector acquisition, and denoising.

The channel estimation procedure relies on the creation of pileups at reference genome positions. As a result, this step considers only bases that are mapped to the reference genome and relies on the CIGAR string information to accurately map bases to reference genome positions. Bases that are designated in the CIGAR string as low-confidence and non-matching (i.e., “soft-clipped”, or simply “clipped”) or inserted relative to the reference genome are not considered during channel estimation. When deletions are indicated in the CIGAR string, reference mapping positions are adjusted accordingly.

The counts vectors are simply histograms of the central base appearing with context strings of length 2*k*. These context strings are unique to the individual and should include bases that are inconsistent relative to the reference genome, since those bases may be true polymorphisms. Thus, during counts vector acquisition the reads retain bases that are marked by the aligner as insertions since those insertions may be inherent to the true sequence. However, as in the channel estimation process, bases that are designated in the CIGAR string as clipped are omitted to avoid large sections of low-confidence base calls. In order to maximize the number of context strings obtained from a processed read, we additionally pad the read with a header and footer of length up to *k* if the read begins or ends, respectively, with bases that are mapped to the reference genome, i.e., not insertions. The padding process, illustrated in Figure 4, allows the denoiser to obtain context information from up to 2*k* additional locations in each read.

During the denoising process, we again require the context string of length 2*k* surrounding a given base. In this step, all bases regardless of their categorization are utilized since bases in the reads that were designated as insertions or deletions relative to the reference genome may very well be true polymorphisms, and soft-clipped regions may benefit from denoising. When the read begins or ends in bases that can be mapped to the reference genome, the read is again padded with bases from the sequence estimate using the same procedure described above in order to maximize the number of bases undergoing denoising.

### Evaluation pipeline workflow

#### Evaluation criteria

To quantify denoiser performance, we used the common performance metrics of true positives (T.P.), false positives (F.P.), and false negatives (F.N.). T.P. variants are the calls present in the gold standard call set, F.P. variants are the calls not present in the gold standard call set, and F.N. variants are those present in the gold standard set but not called. To summarize the changes in T.P., F.P. and F.N. variants identified, we used the following performance metrics: sensitivity (S), which measures the proportion of all the variants that are correctly called (T.P./(T.P. + F.N.)), precision (P), which measures the proportion of called variants that are true (T.P./(T.P. + F.P.)), and F-score (F), which is the harmonic mean of the sensitivity and precision (2(*S* × *P*)/(*S* + *P*)).

#### SAMDUDE parameters

For all denoising experiments, we used a single-sided context length of *k* = 7 (14 bases total in the double-sided context). This context length was chosen for computational feasibility, but also maximizes the number of counts in each context histogram without skewing the histograms towards a uniform distribution, which occurs when *k* is either too small or too large (see Supplementary Tables 4 and 5). For sequence and channel estimation we used a majority threshold of *t*_*m*_ = 0.9 for high confidence in our estimate of the “true” genomic sequence, and also to definitively eliminate potentially confounding effects at heterozygous genomic positions which might not have a clear majority base. Finally, based on experiments with different quality value thresholds (see Supplementary Tables 5 and 6), we attempted denoising only at bases for which the sequencer’s confidence probability *p*, Equation (4)) is less than a chosen confidence threshold *t*_*p*_ = 0.9.

#### Denoising and variant calling pipeline

Individual chromosomes were extracted in binary SAM (BAM) file format from the aligned data sets and sorted using the SAMtools utility^26^. For Musket and RACER denoising, copies of the sorted BAM file were converted from the BAM to FASTQ format via the biobambam2 BAM file processing toolkit^27^. BAM files for each chromosome were also converted to the SAM format. The extracted SAM files were denoised using SAMDUDE.

The denoised FASTQ files were then aligned to a reference file using BWA-mem^28^, generating denoised SAM files. All denoised SAM files then underwent SNP calling using the SNP calling pipeline recommended by the Broad Institute^29–31^, and compared to the gold standard call set. We report results for both raw variants and variants filtered under the GATK Best Practices-recommended variant filtering process. For more details regarding the variant calling, filtering and evaluation pipelines, we refer the reader to the Variant calling pipeline section in the Supplementary data.

#### Computational requirements and machine specifications

We ran most experiments on a workstation computer with 12 Intel Xeon cores at 3.4 GHz and 32 GB of RAM, running Linux Ubuntu 14.04.4. SAMDUDE denoising for the chromosome 11 file of data set 3 was run on a different workstation with 80 Intel Deon cores at 2.2 GHz and 504 GB RAM, running CentOS 7.4.1708. Time and peak computational memory requirements for denoising data sets 1, 2 and 3 using SAMDUDE, Musket and RACER are summarized in the Supplementary Table 7. In its current manifestation, SAMDUDE generally uses about an order of magnitude more memory than Musket and RACER. This is due to the large number of context histogram vectors that SAMDUDE acquires. SAMDUDE also generally requires about an order of magnitude more runtime than Musket and RACER. This result is not surprising, given that SAMDUDE is currently implemented in Python with no parallelization.

## Acknowledgements

The authors would like to thank Jiantao Jiao and Dmitri Pavlichin for helpful discussions and suggestions

## Author contributions statement

I.O., M.H. and T.W. conceived the experiments, I.F. conducted the experiment and analyzed the results. All authors reviewed the manuscript.

## References

1. Costa, F. F. Big data in biomedicine. Drug discovery today 19, 433–440 (2014).

2. Minoche, A. E., Dohm, J. C. & Himmelbauer, H. Evaluation of genomic high-throughput sequencing data generated on illumina hiseq and genome analyzer systems. Genome biology 12, R112 (2011).

3. Boycott, K. M., Vanstone, M. R., Bulman, D. E. & MacKenzie, A. E. Rare-disease genetics in the era of next-generation sequencing: discovery to translation. Nat. Rev. Genet. 14, 681–691 (2013).

4. Laehnemann, D., Borkhardt, A. & McHardy, A. C. Denoising dna deep sequencing data—high-throughput sequencing errors and their correction. Briefings bioinformatics 17, 154–179 (2016).

5. Koh, P. W., Pierson, E. & Kundaje, A. Denoising genome-wide histone chip-seq with convolutional neural networks. Bioinformatics 33, i225–i233, doi: 10.1093/bioinformatics/btx243 (2017). /oup/backfile/content_public/journal/bioinformatics/33/14/10.1093_bioinformatics_btx243/2/btx243.pdf.

6. Molnar, M. & Ilie, L. Correcting illumina data. Briefings bioinformatics 16, 588–599 (2014).

7. Heydari, M., Miclotte, G., Demeester, P., Van de Peer, Y. & Fostier, J. Evaluation of the impact of illumina error correction tools on de novo genome assembly. BMC bioinformatics 18, 374 (2017).

8. Lee, B., Moon, T., Yoon, S. & Weissman, T. Dude-seq: Fast, flexible, and robust denoising for targeted amplicon sequencing. PloS one 12, e0181463 (2017).

9. Ochoa, I., Hernaez, M., Goldfeder, R., Weissman, T. & Ashley, E. Effect of lossy compression of quality scores on variant calling. Briefings bioinformatics 18, 183–194 (2016).

10. Liu, Y., Schröder, J. & Schmidt, B. Musket: a multistage k-mer spectrum-based error corrector for illumina sequence data. Bioinformatics 29, 308–315 (2013).

11. Ilie, L. & Molnar, M. Racer: rapid and accurate correction of errors in reads. Bioinformatics 29, 2490–2493 (2013).

12. Melsted, P. & Pritchard, J. K. Efficient counting of k-mers in dna sequences using a bloom filter. BMC Bioinforma. 12, 333, doi: 10.1186/1471-2105-12-333 (2011).

13. Song, L., Florea, L. & Langmead, B. Lighter: fast and memory-efficient sequencing error correction without counting. Genome Biol. 15, 509, doi: 10.1186/s13059-014-0509-9 (2014).

14. Heo, Y., Ramachandran, A., Hwu, W.-M., Ma, J. & Chen, D. Bless 2: accurate, memory-efficient and fast error correction method. Bioinformatics 32, 2369–2371, doi: 10.1093/bioinformatics/btw146 (2016).

15. Weissman, T., Ordentlich, E., Seroussi, G., Verdú, S. & Weinberger, M. J. Universal discrete denoising: Known channel. IEEE Transactions on Inf. Theory 51, 5–28 (2005).

16. Zook, J. M. et al. Integrating human sequence data sets provides a resource of benchmark snp and indel genotype calls. Nat. biotechnology 32, 246–251 (2014).

17. Cánovas, R., Moffat, A. & Turpin, A. Lossy compression of quality scores in genomic data. Bioinformatics 30, 2130– 2136, doi: 10.1093/bioinformatics/btu183 (2014). /oup/backfile/content_public/journal/bioinformatics/30/15/10.1093_bioinformatics_btu183/2/btu183.pdf.

18. Malysa, G. et al. Qvz: lossy compression of quality values. Bioinformatics 31, 3122–3129, doi: 10.1093/bioinformatics/btv330 (2015). /oup/backfile/content_public/journal/bioinformatics/31/19/10.1093_bioinformatics_btv330/3/btv330.pdf.

19. Gilissen, C., Hoischen, A., Brunner, H. G. & Veltman, J. A. Disease gene identification strategies for exome sequencing. Eur. J. Hum. Genet. 20, 490 (2012).

20. Rabbani, B., Mahdieh, N., Hosomichi, K., Nakaoka, H. & Inoue, I. Next-generation sequencing: impact of exome sequencing in characterizing mendelian disorders. J. human genetics 57, 621 (2012).

21. Bastarache, L. et al. Phenotype risk scores identify patients with unrecognized mendelian disease patterns. Science 359, 1233–1239 (2018).

22. Goldfeder, R. L. et al. Medical implications of technical accuracy in genome sequencing. Genome medicine 8, 24 (2016).

23. Dewey, F. E. et al. Clinical interpretation and implications of whole-genome sequencing. Jama 311, 1035–1045 (2014).

24. Altman, R. B. et al. A research roadmap for next-generation sequencing informatics. Sci. translational medicine 8, 335ps10–335ps10 (2016).

25. Illumina. Reducing whole-genome data storage footprint (white paper, available at https://www.illumina.com/documents/products/whitepapers/whitepaper{_}datacompression.pdf (2014).

26. Li, H. et al. The sequence alignment/map format and samtools. Bioinformatics 25, 2078–2079 (2009).

27. Tischler, G. & Leonard, S. biobambam: tools for read pair collation based algorithms on bam files. Source Code for Biol. Medicine 9, 13 (2014).

28. Li, H. Aligning sequence reads, clone sequences and assembly contigs with bwa-mem. arXiv preprint arXiv:1303.3997 (2013).

29. McKenna, A. et al. The genome analysis toolkit: a mapreduce framework for analyzing next-generation dna sequencing data. Genome research 20, 1297–1303 (2010).

30. DePristo, M. A. et al. A framework for variation discovery and genotyping using next-generation dna sequencing data. Nat. genetics 43, 491–498 (2011).

31. Van der Auwera, G. A. et al. From fastq data to high-confidence variant calls: the genome analysis toolkit best practices pipeline. Curr. protocols bioinformatics 11–10 (2013).

