## Supplementary Data for "Denoising of Aligned Genomic Data"

### Supplementary text

#### SAMDUDE operation

In this section, we describe the operational and parameter choices for SAMDUDE: quality score bin limits, single-sided context length $k$, and confidence probability threshold $t_p$. We also discuss the sequence data coverage requirements for SAMDUDE, and provide details of the random noise addition process.

##### Variantions on SAMDUDE and number of bases denoised

Two "variations" on SAMDUDE were presented in the "Human chromosome denoising with SAMDUDE" subsection of the Results section in the main text: partial denoising and random noise. The results of these variations are summarized in Supplementary Table 1. Supplementary Table 2 lists the total number of bases in the original SAM files, as well as the percent of base changes under each denoiser in the chromosome files for each data set.

##### Quality score binning

In order to reduce the computational burden of estimating and using the channel matrix estimate $\hat{\mathbf{\Pi}}$, we binned the quality scores according to the bin limits in Table 3.

#### Choice of $k$

For discrete universal denoising using DUDE, the optimal single-sided context length $k_n$ depends on both sequence length and alphabet size:

$$k_n = \lceil c \log_{\mathrm{M}} n \rceil$$

with $c < \frac{1}{2}$, noise-free sequence alphabet size M and noise-free sequence length $n$. In the genomic sequencing case, M$=4$, and $n$ ranges from approximately $51 \times 10^6$ up to $249 \times 10^6$. For these values, $k_n = 5$ or 6.

Supplementary Table 4 shows the results of denoising extracted SAM files for chromosomes 11 and 20 NA12878 paired-end WGS data set ERR262997 using $k = 5$ and 6, and confidence probability threshold $t_p = 0.9$. These values of k resulted in improvements in both S and P, but very little change in F-Score. We hypothesized that SAMDUDE's performance could be improved by limiting denoising decisions to local information. We tested larger values of $k$ which should ensure that context counts are taken from the same pileup, while still allowing information to be obtained from misaligned or poorly-mapped reads found elsewhere in the SAM file. Here, we show results for $k = 10$ for chromosome 20. Although the best denoising performance was obtained for $k = 10$, the number of context vectors required used a computationally prohibitive amount of memory. Thus, the denoising results in the main text were obtained using $k = 7$.

#### Choice of confidence probability threshold $t_p$

The choice of confidence probability threshold is a tradeoff. Too low a threshold might prevent numerous noisy bases from being denoised, while too high a threshold might result in changes to bases with very high confidence calls. Supplementary Table 5 shows the results for changes in variant calls with a very high confidence probability threshold of 0.99, corresponding to a quality score of 20. Furthermore, histograms of the original quality scores of bases changed under SAMDUDE for chromosome 20 data sets shows that SAMDUDE-based denoising chooses a different base primarily for bases with a small original quality score (Supplementary Figure 7). Based on these results, we chose to use a lower confidence probability threshold of $t_p = 0.9$ for our denoising tests. This threshold has the additional benefit of reducing computational runtime and preventing over-processing of the original data.

##### Coverage of input data

Supplementary Table 6 shows the results of denoising using SAMDUDE for various values of $k$ on SAM files extracted from NA12878 paired-end WGS data set ERR174324 with 15×-coverage. The lack of improvement, and slight negative effect of SAMDUDE denoising on these lower coverage data are not entirely surprising. SAMDUDE makes significant use of alignment information in order to estimate the noise channel, create counts vectors and to perform denoising, so we expect better denoising performance with higher coverage. Based on these results, we focused our efforts on denoising data with 30× or higher coverage. This tends to be the coverage range of WGS data used in practice, especially for clinical purposes.

#### Operation of state-of-the-art denoising software

In this section we provide the commands and parameters used for denoising with Musket, RACER, BFCounter and Lighter.

##### Musket

Musket was invoked using all default parameters.

Version: 1.1

Website: http://musket.sourceforge.net/homepage.htm

Command:

```
$ musket -p 16 read_1.fastq read_2.fastq -omulti denoised -indorder
```

##### RACER

RACER was invoked using estimated sequence lengths of $63 \times 10^6$ and $135 \times 10^6$ for chromosomes 20 and 11, respectively, and $4 \times 10^9$ as an estimate of the whole human genome length.

Version: 1.0.1

Website: http://www.csd.uwo.ca/ ilie/RACER/

Command:

```
$ racer read_1.fastq read_2.fastq 3000000000
```

##### BFCounter

BFCounter was invoked using estimated sequence lengths of $64 \times 10^6$ and $100 \times 10^6$ for chromosomes 20 and 11, respectively, and $4 \times 10^9$ as an estimate of the whole human genome length.

Version: r181

Website: https://github.com/lh3/bfc

Command:

```
$ bfc -s 100m read_1.fastq.gz read_2.fastq.gz
```

##### Lighter

Lighter was invoked using estimated sequence lengths of $63 \times 10^6$ and $135 \times 10^6$ for chromosomes 20 and 11, respectively, and $4 \times 10^9$ as an estimate of the whole human genome length, and $k$-mer length of 17.

Version: 1.1.1

Website: https://github.com/mourisl/Lighter/

Command:

```
$ lighter read_1.fastq read_2.fastq -K 17 63500000
```

#### Variant calling pipeline

In this section we describe the steps and pipelines used to analyze the effect of base denoising and quality score updating on variant calling.

##### Preprocessing

The first steps in the pipeline are preprocessing steps which include conversion of the SAM file to the BAM format, file sorting, duplicate reads marking, group name adding, file indexing and quality score recalibration.

If the denoised file is in FASTQ format, one extra step of alignment to a reference genome must be performed before the other preprocessing steps. This was done using the BWA mem alignment program and NCBI build 37 of the human reference [`http://www.ncbi.nlm.nih.gov/assembly/GCF$\_$000001405.13/`] as the reference genome. We have included the -M option in *bwa mem* for compatibility with Picard tools, and use the -t option to specify the desired number of threads used for computation.

```
$ bwa mem -t num_threads -M ref.fa read_1.fastq read_2.fastq > aln.sam
```

The denoised reads are now in the SAM file format. All denoised SAM files then underwent SNP calling using the SNP calling pipeline recommended by the Broad Institute [1, 2, 3], which is the following.

The SAM file is converted to the BAM format using samtools, specifying the number of threads with the -@ option and inclusion of the SAM header with the -h option.

```
$ samtools view -@ num_threads -b -h aln.sam > aln.bam
```

The BAM file is then sorted using samtools, with a temporary file prefix specified with the -T option, and the output file format specified with the -O option.

```
$ samtools sort -T ./tmp -@ num_threads -O bam aln.bam > aln.sorted.bam
```

Duplicates are marked using Picard tools [1], with M specifying the file to which metrics calculated during the duplication marking process are written. Note that marking the duplicates is sufficient to exclude them from downstream processes.

```
$ java -jar picard.jar MarkDuplicates I=aln.sorted.bam \
      O=aln.sorted.dedup.bam M=metrics.txt ASSUME_SORTED=true
```

Read group names are then added to the file, with read group parameters specified by RGID, RGLB, RGPL, RGPU and RGSM.

```
$ java -jar picard.jar AddOrReplaceReadGroups INPUT=aln.sorted.dedup.bam \
      OUTPUT=aln.sorted.dedup.rg.bam RGID=group1 RGLB=lib1 \
      RGPL=illumina RGPU=unit1 RGSM=NA12878
```

Then, the BAM file is indexed.

```
$ java -jar picard.jar BuildBamIndex I=$aln.sorted.dedup.rg.BAM
```

Finally, the quality scores are recalibrated using the Genome Analysis Toolkit [1] Base Quality Score Recalibration workflow, and the NCBI build 37 of the human reference as the reference genome.

```
$ java -jar GenomeAnalysisTK.jar -nct num_threads -T BaseRecalibrator -R pa
      -I aln.sorted.dedup.rg.bam \
      -knownSites bundle_2.8/dbsnp_138.b37.vcf \
      -knownSites bundle_2.8/Mills_and_1000G_gold_standard.indels.b37.vcf \
      -knownSites bundle_2.8/1000G_phase1.indels.b37.vcf -o bqsr.data
```

```
$ java -jar GenomeAnalysisTK.jar -nct num_threads -T PrintReads -R pathHuma
      -I aln.sorted.dedup.rg.bam -BQSR recal_data -o aln.sorted.dedup.rg.re
```

##### Variant calling and filtering

We use the GATK Haplotype Caller for variant calling, specifying the target chromosome with the -L option.

```
$ java -jar GenomeAnalysisTK.jar -T HaplotypeCaller -R pathHumanReference \
      -I aln.sorted.dedup.rg.recal.bam -L targetRegion \
      --genotyping_mode DISCOVERY -stand_emit_conf 10 -stand_call_conf 30 -
```

Finally, we use the Illumina open source hapolotype comparison tool hap.py (https://github.com/Illumina/hap.py#happy) to compare the variant calls of the denoised files to those of the original file, for both raw variant calls as well as those filtered by the GATK Best Practices variant filtering procedure [2]. The hap.py evaluation pipeline is also used to filter variants and extract true and false positive values.

```
$ python hap.py ground_truth.vcf $raw_VCF -f ground_truth.bed -o results -r
$ python rep.py -o results.html -l tsv_file
```

#### Data sets

The data sets used in this study were obtained from the following sources: ERR262997 (data set 1) with 30×-coverage (http://www.ebi.ac.uk/ena/data/view/ERA207860), CEUTrio.HiSeq.WGS (data set 2) with 100×-coverage (ftp://gsapubftp-anonymous@ftp.broadinstitute.org/bundle/b37/CEUTrio.HiSeq.WGS.b37.NA12878.bam), and NA12878_V2.5_Robot_2 (data set 3) with 40×-coverage (https://www.garvan.org.au/research/kinghorn-centre-for-clinical-genomics/clinical-genomics/sequencing-services/sample-data).

#### Computing requirements

Time and peak computational memory requirements for denoising data sets 1, 2 and 3 using SAMDUDE, Musket, RACER, BFCounter and Lighter are summarized in Supplementary Table 7. Thw reported Times the time necessary to process 1 MB of reads in the original file (SAM for SAMDUDE, and FASTQ for Musket, RACER, BFCounter and Lighter). In its current manifestation, SAMDUDE generally uses about an order of magnitude more memory than Musket and RACER, and nearly two orders of magnitude more memory than BFCounter and Lighter. This is due to the large number of context histogram vectors that SAMDUDE acquires. SAMDUDE also generally requires about one to two orders of magnitude more runtime than the state-of-the-art denoisers. This result is not surprising, given that SAMDUDE is currently implemented in Python with no parallelization.

#### Sensitivity vs. precision curves

Supplementary Figures 1, 2, 3, 4, 5, and 6 show precision as a function of sensitivity for variant call sets filtered by QD. The odd-numbered figures include an additional rightmost point corresponding to the raw variant calls corresponding to those reported in all other tables and figures, while the even-numbered figures omit the rightmost point for ease of visualization.

In all curves, the performance of SAMDUDE is compared with lossy quality score compressors P-Block and R-Block, as well as with Musket and RACER. For ease of visualization, the results of denoising with BFCounter and Lighter are omitted.

#### Tables and Figures

| | data set | SAMDUDE ΔS [%] | SAMDUDE ΔP [%] | SAMDUDE ΔF [%] | Partial denoising ΔS [%] | Partial denoising ΔP [%] | Partial denoising ΔF [%] | Random noise ΔS [%] | Random noise ΔP [%] | Random noise ΔF [%] |
|---|---|---|---|---|---|---|---|---|---|---|
| raw | 1 | 0.04 | 0.12 | 0.08 | -0.01 | 0.19 | 0.09 | -1.39 | 1.58 | 0.10 |
| | 2 | – | 0.95 | 0.47 | 0.01 | 0.05 | 0.03 | -0.88 | 1.00 | 0.06 |
| | 3 | – | 0.02 | 0.01 | – | -0.01 | – | – | 0.03 | 0.01 |
| GATK filtered | 1 | 0.03 | 0.11 | 0.07 | -0.01 | 0.12 | 0.06 | -2.29 | 0.78 | -0.76 |
| | 2 | -0.01 | 0.75 | 0.37 | 0.01 | 0.02 | 0.02 | -1.39 | 0.80 | -0.29 |
| | 3 | 0.01 | – | – | – | 0.01 | 0.01 | -0.01 | 0.02 | 0.01 |

Supplementary Table 1: Changes (Δ) in sensitivity (S), precision (P) and F-score (F) under SAMDUDE, SAMDUDE-denoised reads with original quality scores (Partial denoising), and random noise (Random noise) calculated relative to the original file. Positive Δ indicates improvement with respect to the original data, and horizontal lines indicate no change.

| chr | data set | $n$ | SAMDUDE [%] | Musket [%] | RACER [%] | BFCounter [%] | Lighter [%] |
|---|---|---|---|---|---|---|---|
| 11 | 1 | 5,806,522,969 | 0.36 | 0.34 | 8.41 | 0.23 | 0.42 |
| | 2 | 11,960,009,536 | 1.80 | 0.53 | 1.26 | 0.55 | 0.62 |
| | 3 | 6,769,559,684 | 0.07 | 0.80 | 1.84 | 0.89 | 0.62 |
| 20 | 1 | 2,538,750,907 | 0.35 | 0.30 | 8.76 | 0.26 | 0.42 |
| | 2 | 5,206,460,817 | 1.80 | 0.59 | 1.34 | 0.63 | 0.73 |
| | 3 | 3,064,700,879 | 0.11 | 1.77 | 1.99 | 0.99 | 0.70 |

Supplementary Table 2: Total number of bases in the original SAM files ($n$) compared to the percentage of base changes recommended under the five denoisers.

| Quality score bin | Quality score range |
|---|---|
| 1 | $< 2$ |
| 2 | 2–9 |
| 3 | 10–19 |
| 4 | 20–24 |
| 5 | 25–29 |
| 6 | 30–34 |
| 7 | 35–39 |
| 8 | $\geq 40$ |

Supplementary Table 3: Quality score bin labels and ranges.

| | | Raw variant calls | | | | GATK filtered variant calls | | | |
|---|---|---|---|---|---|---|---|---|---|
| chr | $k$ | $\Delta$C | $\Delta$S [%] | $\Delta$P [%] | $\Delta$F [%] | $\Delta$C | $\Delta$S [%] | $\Delta$P [%] | $\Delta$F [%] |
| 11 | 5 | 26 | 0.01 | 0.03 | – | 50 | 0.01 | 0.02 | – |
| | 6 | -199 | 0.03 | 0.17 | 0.10 | -80 | 0.03 | 0.11 | 0.10 |
| 20 | 5 | 59 | 0.04 | 0.07 | – | 59 | 0.04 | 0.06 | – |
| | 6 | -19 | 0.01 | 0.10 | – | 18 | 0.01 | 0.05 | – |
| | 10 | 373 | 0.21 | 0.16 | 0.20 | 72 | 0.20 | 0.12 | 0.10 |

Supplementary Table 4: Results of denoising data set ERR262997 with different values of $k$, with changes ($\Delta$) in T.P. and F.P. calculated relative to the original file. C is the number of additional variants called for each condition relative to the variant call set for the original file. For S, P and F, positive $\Delta$ indicates improvement with respect to the original data, and horizontal lines indicate no change.

| | | Raw variant calls | | | | GATK filtered variant calls | | | |
|---|---|---|---|---|---|---|---|---|---|
| chr | $k$ | ΔC | ΔS [%] | ΔP [%] | ΔF [%] | ΔC | ΔS [%] | ΔP [%] | ΔF [%] |
| 11 | 5 | 71 | 0.01 | 0.03 | – | 103 | 0.01 | 0.02 | – |
| | 6 | -238 | – | 0.20 | 0.10 | -112 | – | 0.12 | 0.10 |
| 20 | 5 | 79 | 0.05 | 0.05 | – | 92 | 0.05 | 0.03 | |
| | 6 | -23 | -0.01 | 0.10 | – | 20 | -0.03 | 0.03 | – |

Supplementary Table 5: Results of denoising data set ERR262997 with various $k$ with confidence threshold $t_p = 0.99$. C is the number of additional variants called for each condition relative to the variant call set for the original file, and changes (Δ) were calculated relative to the original file. Positive Δ indicates improvement with respect to the original data, and horizontal lines indicate no change.

| | | Raw variant calls | | | | GATK filtered variant calls | | | |
|---|---|---|---|---|---|---|---|---|---|
| chr | $k$ | ΔC | ΔS [%] | ΔP [%] | ΔF [%] | ΔC | ΔS [%] | ΔP [%] | ΔF [%] |
| 11 | 5 | -13 | 0.01 | – | – | -7 | 0.01 | – | – |
| | 6 | -209 | – | 0.01 | – | -220 | -0.02 | 0.01 | – |
| | 7 | -451 | -0.03 | 0.03 | – | -382 | -0.03 | – | – |
| 20 | 5 | 18 | – | -0.01 | – | 39 | 0.01 | – | – |
| | 6 | -111 | -0.03 | 0.01 | – | -103 | -0.04 | – | – |
| | 7 | -153 | -0.02 | – | – | -117 | -0.03 | – | – |
| | 10 | 235 | 0.05 | -0.04 | – | 189 | 0.03 | -0.02 | – |

Supplementary Table 6: Results of denoising low coverage data set ERR174324 with confidence threshold $t_p = 0.9$. C is the number of additional variants called for each condition relative to the variant call set for the original file, and changes (Δ) were calculated relative to the original file. Positive Δ indicates improvement with respect to the original data, and horizontal lines indicate no change.

| | | SAMDUDE | | Musket | | RACER | | BFCounter | | Lighter | |
|---|---|---|---|---|---|---|---|---|---|---|---|
| chr | data set | Memory [GB] | Time [s/MB] | Memory [GB] | Time [s/MB] | Memory [GB] | Time [s/MB] | Memory [GB] | Time [s/MB] | Memory [GB] | Time [s/MB] |
| 11 | 1 | 27.45 | 0.89 | 2.76 | 0.42 | 7.66 | 0.05 | 5.18 | 0.55 | 0.66 | 0.17 |
| | 2 | 31.79 | 3.77 | 6.53 | 0.56 | 16.07 | 0.08 | 5.22 | 0.36 | 0.66 | 0.12 |
| | 3 | 77.57 | 4.37 | 5.52 | 0.76 | 11.05 | 0.06 | 5.16 | 0.43 | 0.66 | 0.14 |
| 20 | 1 | 31.90 | 6.12 | 1.06 | 0.60 | 3.50 | 0.06 | 2.94 | 0.37 | 0.33 | 0.17 |
| | 2 | 31.74 | 2.58 | 2.14 | 0.72 | 7.35 | 0.05 | 2.94 | 0.35 | 0.33 | 0.14 |
| | 3 | 31.86 | 4.37 | 1.73 | 0.76 | 5.08 | 0.06 | 2.85 | 0.4 | 0.33 | 0.11 |

Supplementary Table 7: Time and peak memory requirements for denoising individual chromosome files from data sets 1, 2 and 3 using the five denoisers. RACER, BFCounter and Lighter requirements are averaged between the paired-end files.

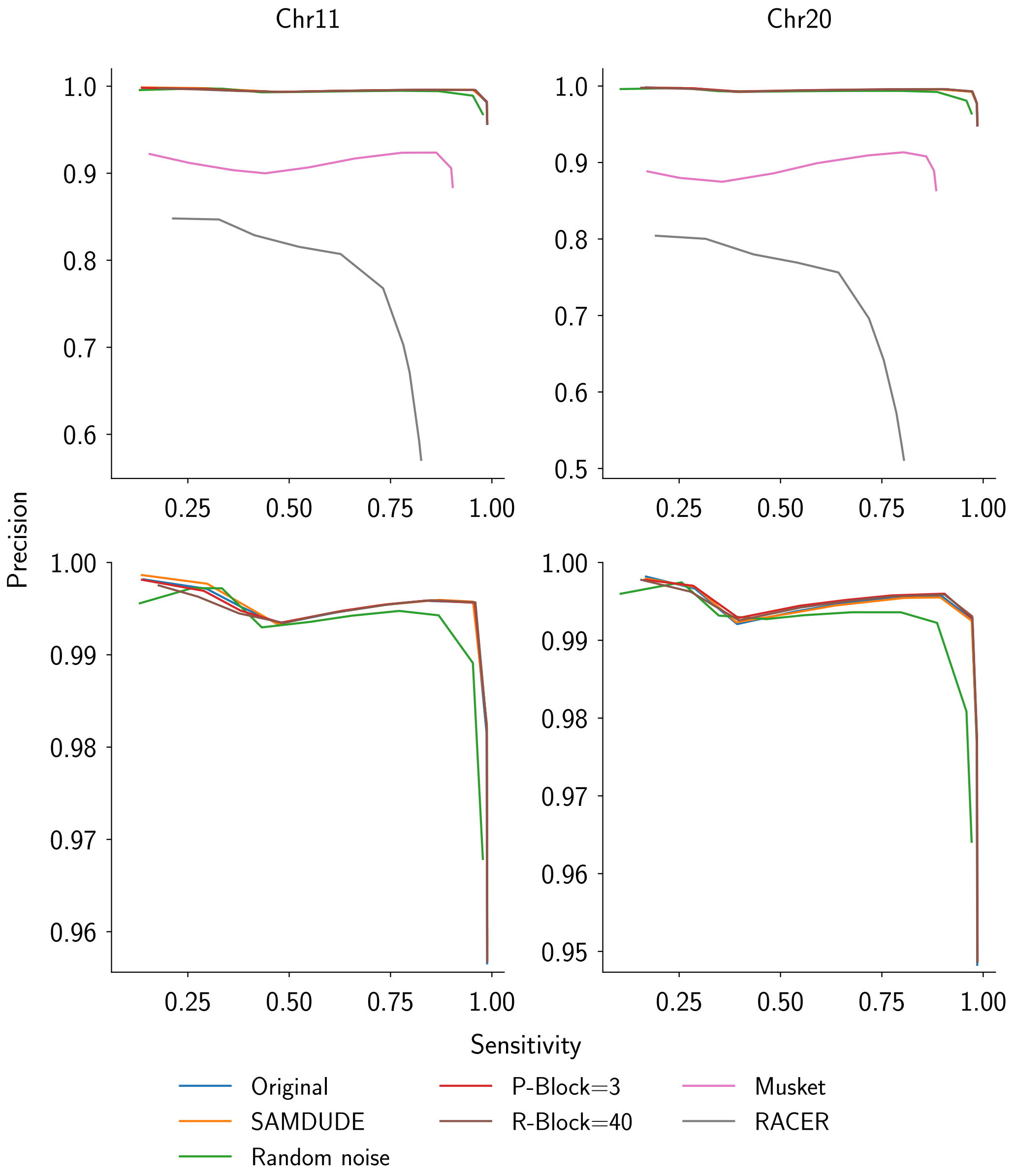


Supplementary Figure 1: Sensitivity vs. precision curves for the data set 1 variant call set filtered at $10^{\text{th}}$ percentiles, starting with no filtering. The top row shows results for all files, and the bottom row omits results for Musket- and RACER-denoised files for closer comparison.

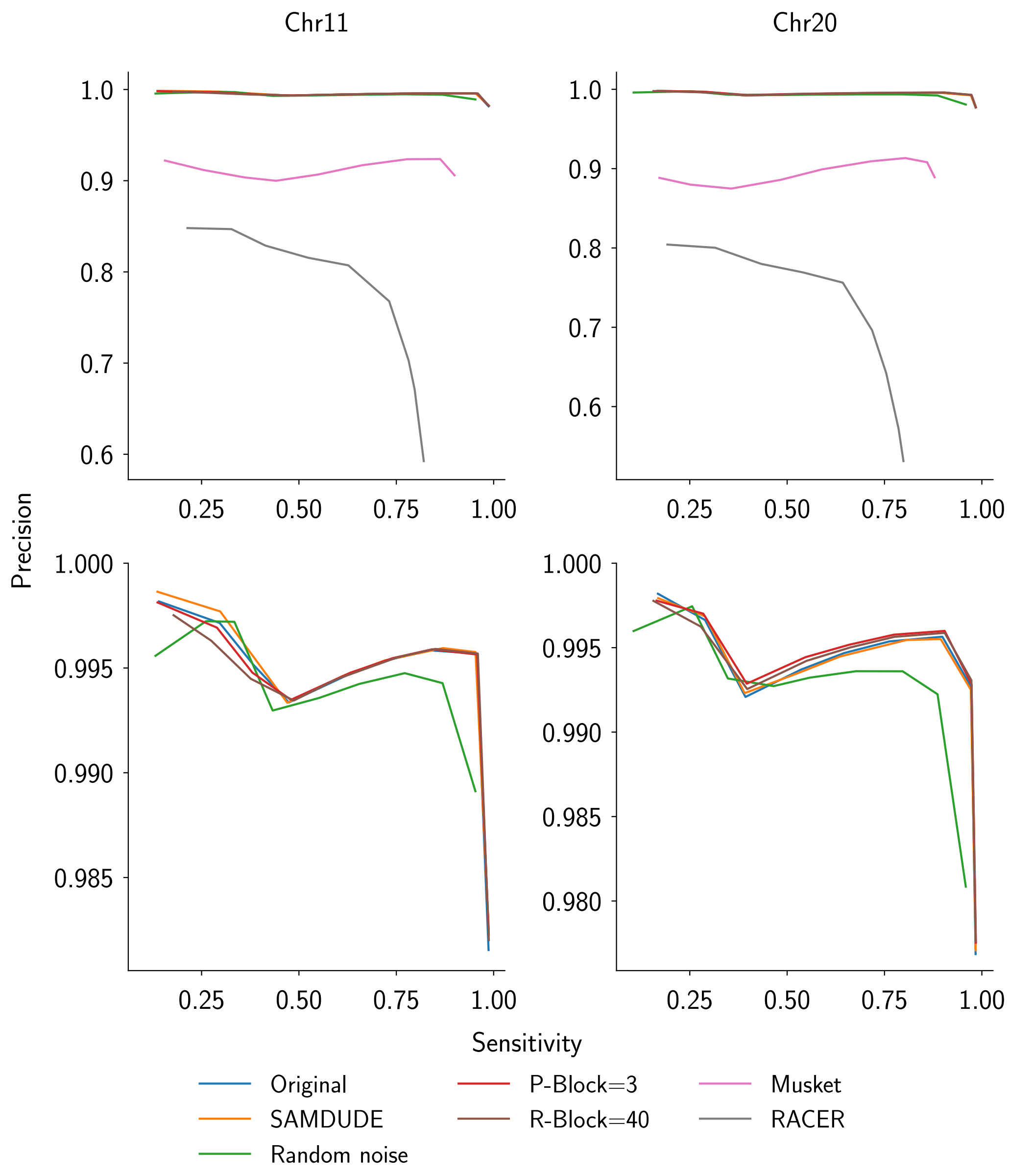


Supplementary Figure 2: The same sensitivity vs. precision curves as in Supplementary Figure 1, but with the rightmost point in each subplot removed for ease of visualization.

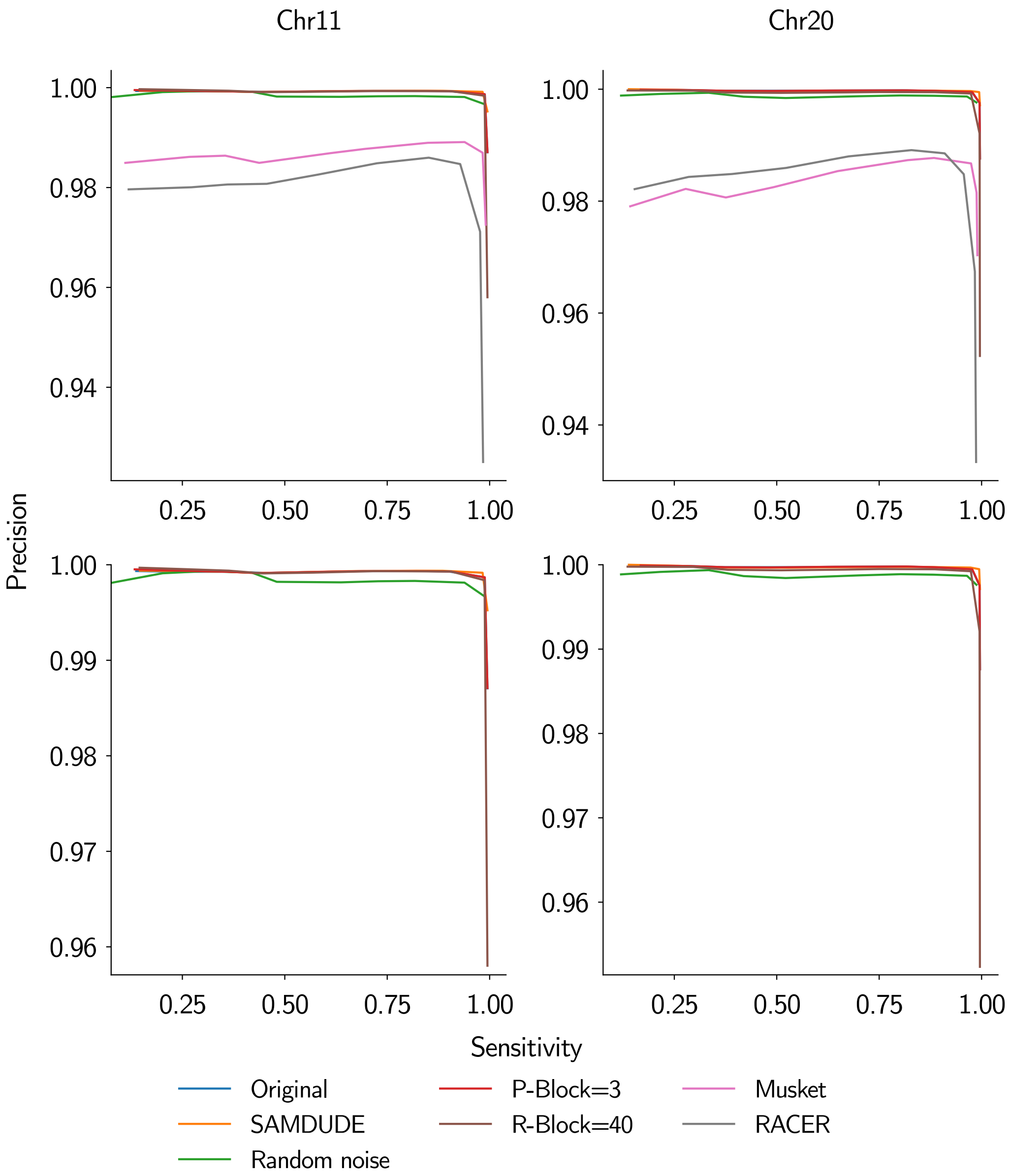


Supplementary Figure 3: Sensitivity vs. precision curves for the data set 2 variant call set filtered at $10^{\text{th}}$ percentiles, starting with no filtering. The top row shows results for all files, and the bottom row omits results for Musket- and RACER-denoised files for closer comparison.

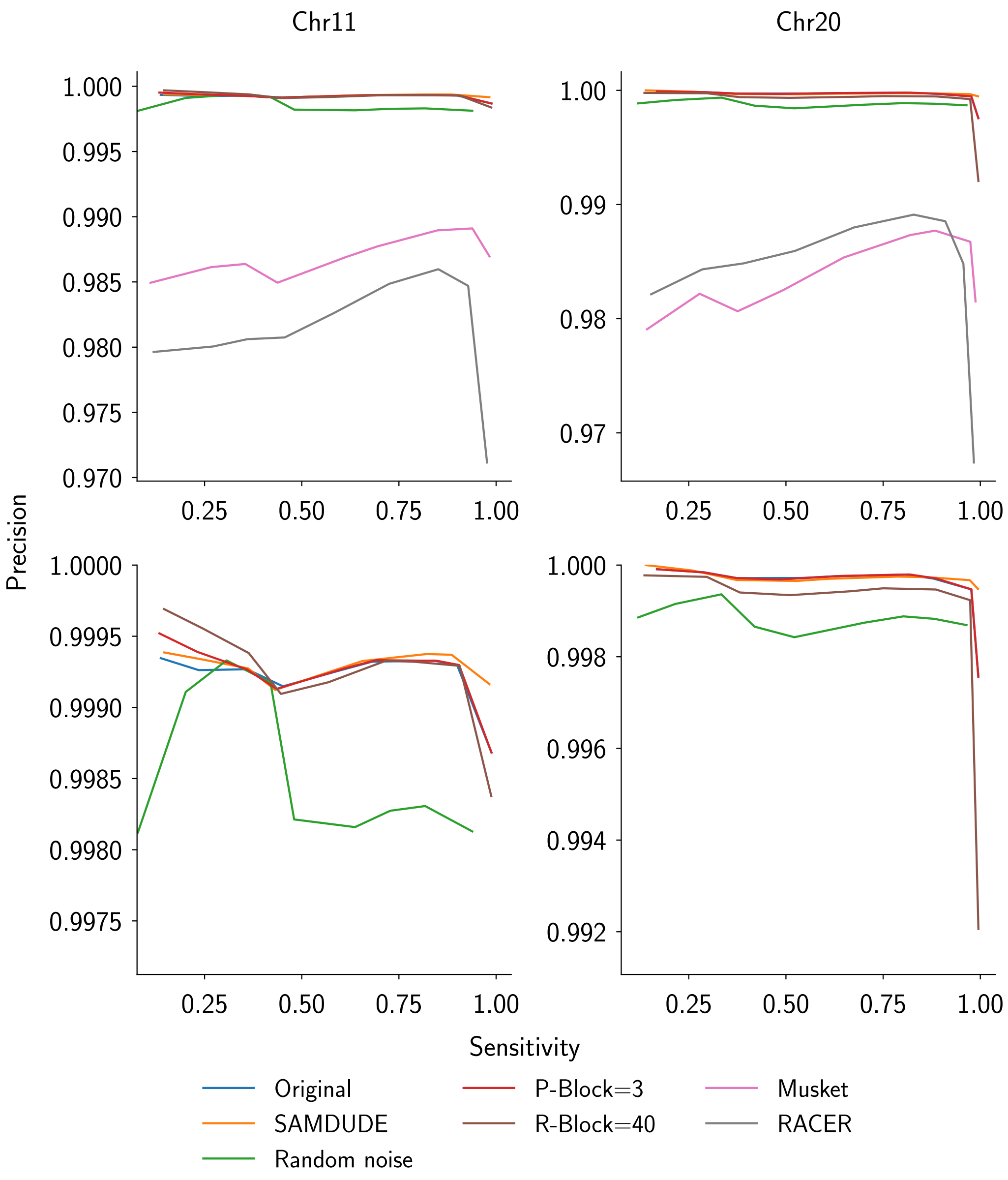


Supplementary Figure 4: The same sensitivity vs. precision curves as in Supplementary Figure 3, but with the rightmost point in each subplot removed for ease of visualization.

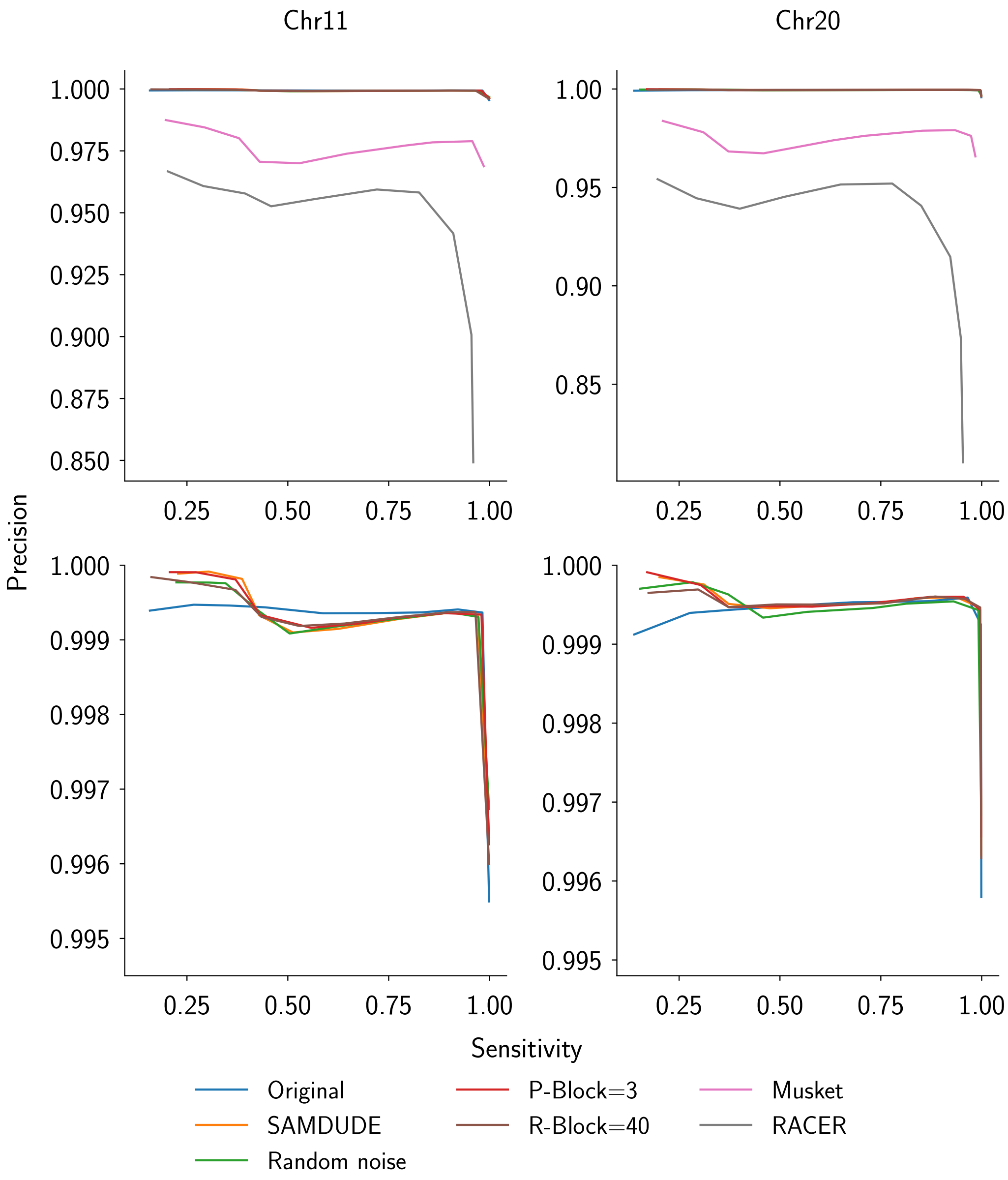


Supplementary Figure 5: Sensitivity vs. precision curves for the data set 3 variant call set filtered at $10^{th}$ percentiles, starting with no filtering. The top row shows results for all files, and the bottom row omits results for Musket- and RACER-denoised files for closer comparison.

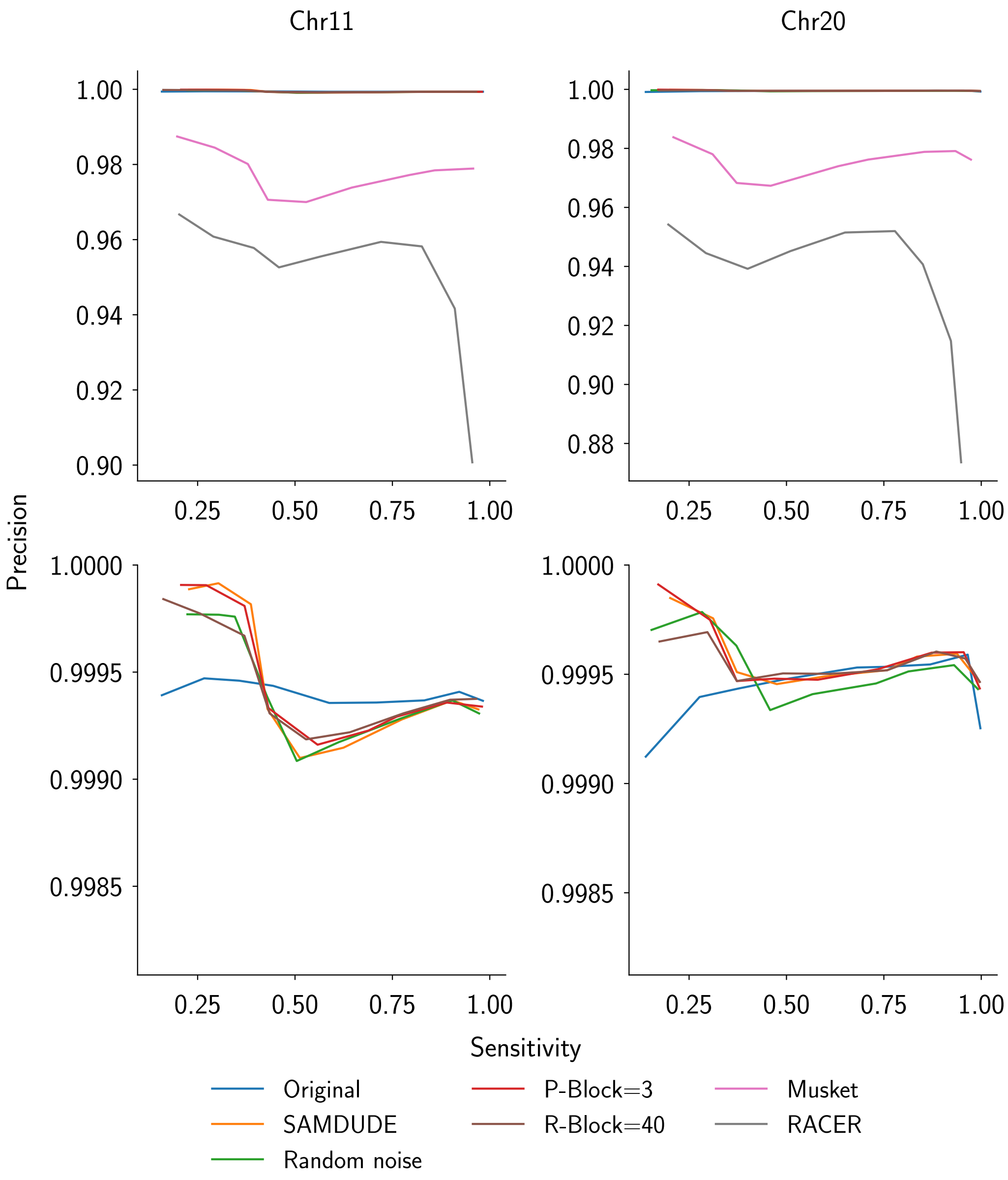


Supplementary Figure 6: The same sensitivity vs. precision curves as in Supplementary Figure 5, but with the rightmost point in each subplot removed for ease of visualization.

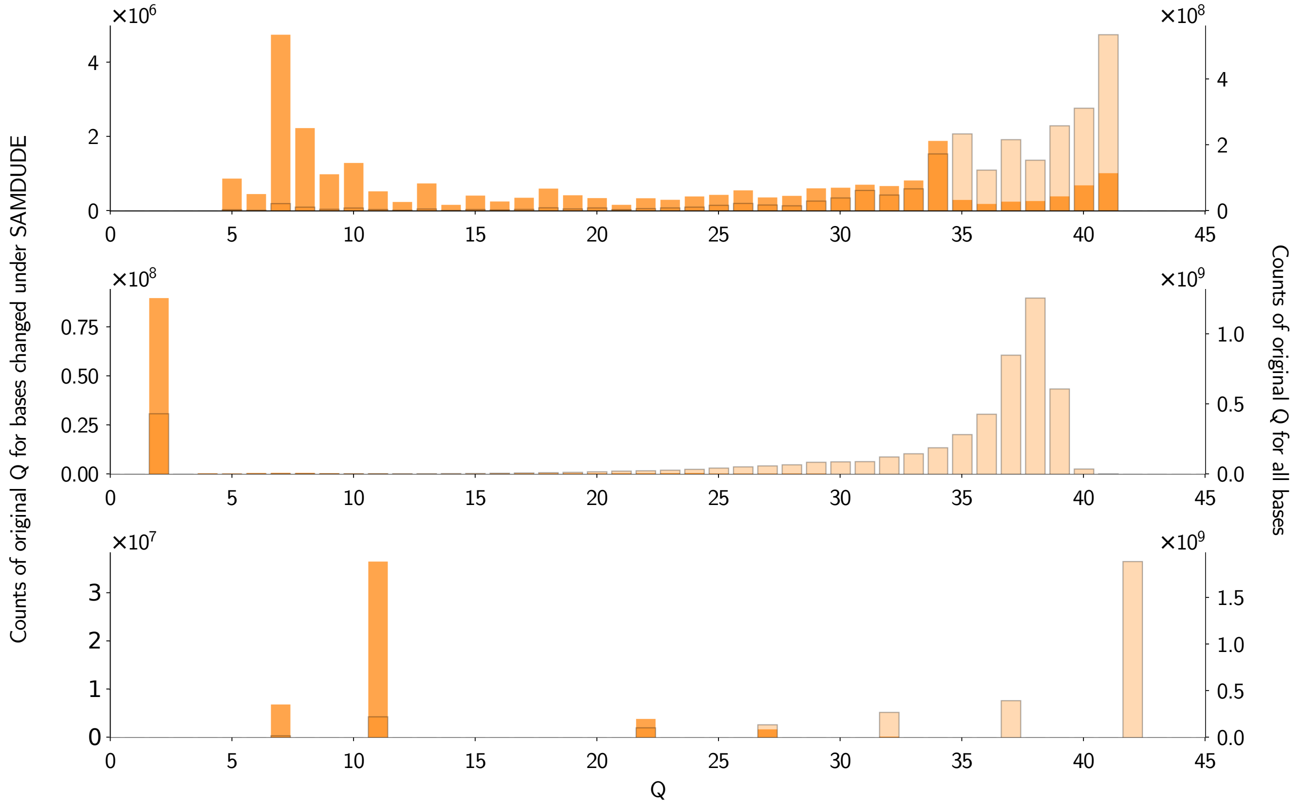


Supplementary Figure 7: Histograms of original quality scores for all bases (light orange, outlined) and for bases changed under the SAMDUDE denoising rule (dark orange) for chromosome 20 SAM files.

### References

[1] A. McKenna, M. Hanna, E. Banks, A. Sivachenko, K. Cibulskis, A. Kernytsky, K. Garimella, D. Altshuler, S. Gabriel, M. Daly *et al.*, "The genome analysis toolkit: a mapreduce framework for analyzing next-generation dna sequencing data," *Genome research*, vol. 20, no. 9, pp. 1297–1303, 2010.

[2] M. A. DePristo, E. Banks, R. Poplin, K. V. Garimella, J. R. Maguire, C. Hartl, A. A. Philippakis, G. Del Angel, M. A. Rivas, M. Hanna *et al.*, "A framework for variation discovery and genotyping using next-generation dna sequencing data," *Nature genetics*, vol. 43, no. 5, pp. 491–498, 2011.

[3] G. A. Van der Auwera, M. O. Carneiro, C. Hartl, R. Poplin, G. del Angel, A. Levy-Moonshine, T. Jordan, K. Shakir, D. Roazen, J. Thibault *et al.*, "From fastq data to high-confidence variant calls: the genome analysis toolkit best practices pipeline," *Current protocols in bioinformatics*, pp. 11–10, 2013.
